# Structure of the Core of the Type Three Secretion System Export Apparatus

**DOI:** 10.1101/249128

**Authors:** Lucas Kuhlen, Patrizia Abrusci, Steven Johnson, Joseph Gault, Justin Deme, Joseph Caesar, Tobias Dietsche, Mehari Tesfazgi Mebrhatu, Tariq Ganief, Boris Macek, Samuel Wagner, Carol V. Robinson, Susan M. Lea

**Affiliations:** Sir William Dunn School of Pathology, University of Oxford, South Parks Road, Oxford OX1 3RE, UK.; Department of Chemistry, University of Oxford, 12 Mansfield Road, Oxford OX1 3TA, UK.; Central Oxford Structural Microscopy and Imaging Centre, University of Oxford, South Parks Road, Oxford OX1 3RE.; Section of Cellular and Molecular Microbiology, Interfaculty Institute of Microbiology and Infection Medicine (IMIT), University of Tübingen, Elfriede-Aulhorn-Str. 6, 72076 Tübingen, Germany; Proteome Center Tübingen, University of Tübingen, Auf der Morgenstelle 15, 72076 Tübingen,Germany; German Center for Infection Research, Partner-site Tübingen, Elfriede-Aulhorn-Str. 6, 72076 Tübingen, Germany

## Abstract

Export of proteins through type three secretion systems is critical for bacterial motility and virulence of many major bacterial pathogens. Three putative integral membrane proteins (FliP/FliQ/FliR) are suggested to form the core of an export gate in the inner membrane, but their structure, assembly and location within the final nanomachine remain unclear. We here present the structure of this complex at 4.2 Å by cryo-electron microscopy. None of the subunits adopt canonical integral membrane protein topologies and common helix-turn-helix structural elements allow them to form a helical assembly with 5:4:1 stoichiometry. Fitting of the structure into reconstructions of intact secretion systems localize the export gate as a core component of the periplasmic portion of the machinery, and cross-linking experiments confirm this observation. This study thereby identifies the export gate as a key element of the secretion channel and implies that it primes the helical architecture of the components assembling downstream.

**One Sentence Summary:** The core of the T3SS export gate forms a supra-membrane helical assembly

---

Type three secretion systems (T3SS) are nanomachines that span the bacterial cell envelope and provide a conduit for protein export from the bacterial cytoplasm ^1^. Two classes of T3SS exist. The first is dedicated to the export and assembly of bacterial flagella. The second, termed the injectisome, allows the delivery of effector proteins directly into the eukaryotic host cell cytoplasm ^2^. Both of these classes are associated with the pathogenicity of a wide range of clinically relevant bacteria ^3^. T3SS are assembled from a basal body consisting of a series of concentric oligomeric protein rings across the inner and outer membranes, from which the helical hook and flagellum or needle structures project ^2,4,5^. Proteins associated with the cytoplasmic face of the basal body select proteins for export that are then transferred to a set of 5 membrane associated proteins roughly located to center of the inner-membrane ring. These components (FliP/FliQ/FliR/FlhB/FlhA in the flagellar system and SctR/SctS/SctT/SctU/SctV in injectisomes, hereafter referred to as P/Q/R/B/A) are collectively termed the export apparatus (EA) and are directly implicated in translocation of substrates across the bacterial envelope ^6-8^.

A combination of many structural techniques has led to atomic models for most of the soluble components, the circularly symmetric rings that compose the majority of the basal body and for both flagellar rod/hook/flagellum and injectisome needle ^4^. However, the EA remains poorly understood, with the topology and number of membrane helices of the three most hydrophobic proteins (**Fig. 1A**, P/Q/R) a subject of debate and with conflicting reports of stoichiometry between flagellar ^7^ and injectisome ^9,10^ T3SS. Given the high levels of structural homology amongst all T3SS structural components revealed to date, and the high levels of sequence conservation in the EA components in particular, it seemed likely that discrepancies between the systems reflected varying experimental approaches rather than true differences in the way in which this core component of the apparatus is assembled. P, Q & R are often encoded within a single operon and we decided to use a combination of biochemistry, native mass spectrometry and cryo-electron miscroscopy to investigate the structure and assembly of these core components in both flagellar and injectisome T3SS.

**Fig. 1.**
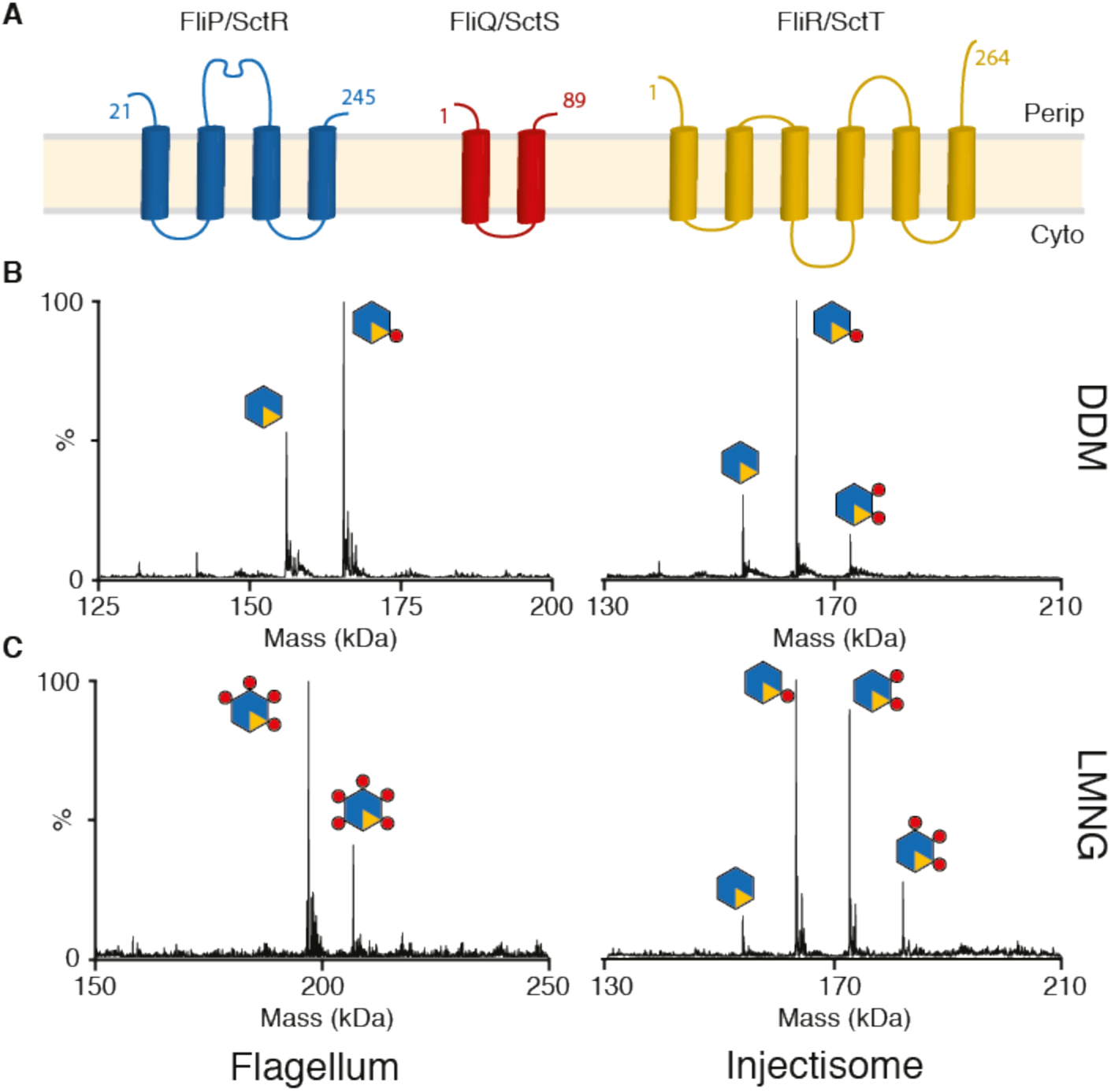
Stoichiometry of the PQR complexes from both Flagellar and Injectisome T3SSs revealed by native mass spectromety. (nMS) **(A)** Consensus topology predictions for the flagellar (FliP, FliQ & FliR) and injectisome (SctR, SctS & SctT) export gate components P, Q & R, numbered according to the *S. enterica* flagellar sequences. The orientation of the termini of Q and R with respect to the membrane has been previously debated and they are shown in here in the same orientation as P. **(B)** Deconvoluted native mass spectra of complexes extracted and purified in DDM reveals a P_5_R_1_ core complex with variable numbers of Q **(C)** Complexes purified in the less harsh detergent LMNG contain more Q subunits, with up to the five copies of Q seen in a flagellar species complex (Table S1).

We previously used native mass spectrometry (nMS) to characterize a P_5_R_1_ complex from the *Salmonella enterica* injectisome system ^9^, whilst others have used negative stain electron microscopy to define a flagellar P_6_ complex ^7^. To attempt to resolve whether flagellar and injectisome T3SS are differently assembled we expressed the P/Q/R components from a range of flagellar and injectisome systems, each from a single operon with a C-terminal dual strep-tag on R for purification. Using the detergent DDM for extraction, high resolution nMS demonstrated that the core of the export apparatus is formed by a P_5_R_1_ complex in all the systems studied (**Fig. 1B, Table S1**). Depending on the species expressed we observed a variable number of Q subunits associated with this core complex (**Fig. 1B**). Dissociating the intact core complex inside the MS using high levels of collisional activation resulted in formation of subcomplexes of all possible combinations of P and R (i.e. P_4_R_1_, P_3_R_1_, P_2_R_1_, P_1_R_1_, P_5_, P_4_, P_3_, P_2_, P_1_), suggesting that the R component is integrated into the complex rather than peripherally associated with a P_5_ ring (**Fig. S1**).

Since we observed variation in the copy number of the Q subunit we investigated different purification strategies and examined the composition of the EA by nMS. Using the less harsh detergent LMNG to solubilize the complex revealed more associated Q subunits, with up to P_5_Q_5_R_1_ complexes being seen (**Fig. 1C, Table S1**). This suggests that multiple Q subunits are loosely associated with a P_5_R_1_ core.

Complexes purified in a range of detergents and amphipols were used to prepare cryo-EM grids with a focus on detergents revealed by nMS to be less destructive. A sample of *S. enterica* FliPQR, purified in LMNG (**Fig. S2**), gave a range of different orientations (**Fig. S3**) and led to a reconstruction of the complex at 4.2Å (**Fig. 2A**). This allowed us to build models for all three components (**Fig. 2B** and Movie S1). The complex is a P_5_Q_4_R_1_ assembly consistent with the dominant species in our earlier nMS data of the flagellar system (**Fig. 1C, Fig. S1, Table S1**): P and R are intimately associated in a pseudo-hexameric, closed structure, with the Q subunits peripherally associated around the outside of the core P_5_R_1_ complex. All three subunits form extended structures, built predominantly from pairs of kinked anti-parallel helices, with a common orientation of N and C termini, in conflict with earlier topology predictions. Many of the conserved charged residues that lead to motility defects when mutated are involved in intra‐ and inter-chain salt bridges in the structure. Each Q is associated with a P, with the Q packed between P and R having a larger buried surface area than the other copies (**Table S2**), implying this is the most tightly bound Q that remains associated in DDM (**Fig. 1B**). This leaves one orphan P unassociated with a copy of Q, and modeling of a fifth Q onto this P-subunit leads to clashes with the neighboring R-subunit. Furthermore, Glu46 in the first 3 Q subunits (Q_1_-Q_3_) bridges to Lys54 in the neighboring Q (Q_2_-Q_4_), while the fourth Glu46 forms a salt bridge with the highly conserved Arg206 in R. All of the interfaces involve residue pairs that have been shown to co-evolve ^11^ with homo-typic interaction surfaces explaining contacts that couldn’t be reconciled in a single protein model. Further investigation reveals that R is a structural fusion of P and Q (**Fig. 2C**), thereby explaining the role of R in bridging between the P_5_ and Q_4_ components of the complex. The major structural difference between a P/Q fusion and the R protein lies in the previously characterized “periplasmic domain” of P ^7^. This domain decorates the outside of the complex, and mapping of sequence conservation onto the surface of the complex reveals this to be the most variable region of the structure (**Fig. 2D**). An earlier crystal structure of this region from *Thermotoga maritima* overlays well with our structure (**Fig. S4**, ^7^). Given the robust prediction that all three subunits are integral membrane proteins, mapping hydrophobicity onto the surface of the model reveals a surprisingly small hydrophobic strip near the base of the structure. Most of the exposed hydrophobicity is contributed by residues of Q, with the majority of the P/R hydrophobics buried in assembly of the complex (**Fig. 2E**). Whilst the surface area buried in assembly of the complex is large for all subunits presumably reflecting strong association (**Table S2**), we note that there are several hydrophobic cavities within the structure which we assume are occupied by buried lipids or detergent that are not resolved at the resolution of our current maps. Based on the location of the detergent belt in our structure it is noteworthy that many of the predicted TM-helices (**Fig. 2F**) are more than 50 Å above this location. Whilst it is possible that following synthesis all predicted TM-helices are membrane localized, folding and assembly must drive removal from the lipid bilayer. The overall packing of the final object demonstrates the difficulties of predicting transmembrane regions in complex multimeric membrane proteins.

**Fig. 2.**
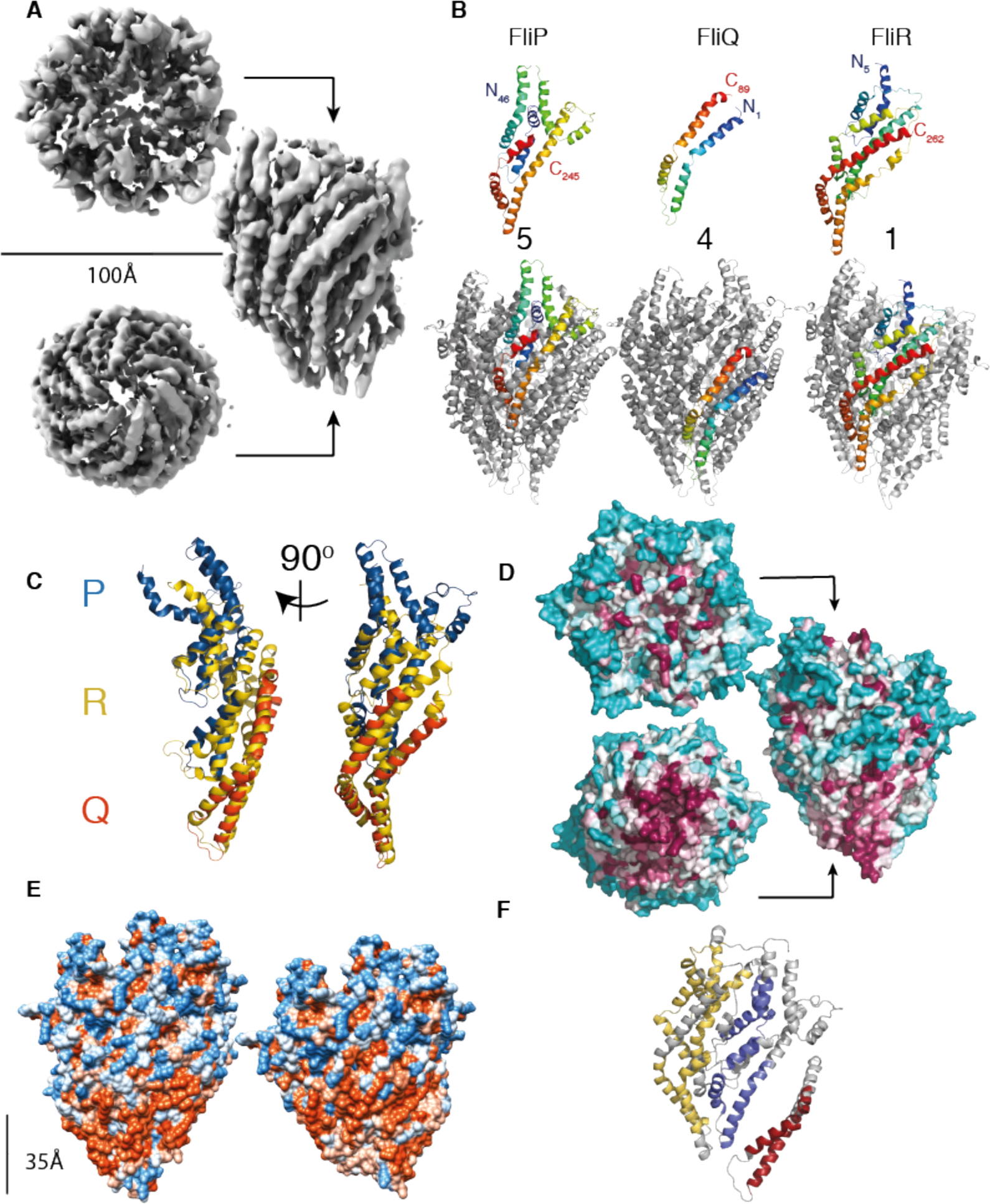
Structure of the flagellar P_5_Q_4_R_1_ complex revealed at 4.2Å by cryo-electron microscopy. **(A)** Cryo-EM map of the P5Q4R1 complex reconstructed from 98000 particles with Cl symmetry. The complex is ~120 Å in height and the top has a diameter of ~100 Å. **(B)** Structures of the monomeric chains and location of an example of each within the full assembly. Each monomer is colored from blue to red at the N and C termini respectively. **(C)** Superposition of a P (blue):Q (orange) heterodimer onto R (yellow) reveals that R is a fusion of the two shorter proteins. **(D)** Analysis of conservation using the CONSURF server^25^ reveals that the bottom of the complex is the major conserved external surface, while the region of P that adorns the outside is highly variable. The relative degree of conservation is colored dark purple for highly conserved to cyan for variable residues. **(E)** The dimensions of the complex are incompatible with all of the proteins being membrane embedded. Mapping hydrophobicity onto the surface of the intact complex (LHS), and the complex of just FliP_5_R_1_ (RHS), reveals that only the middle portion of FliQ is likely to interact with a membrane environment in the assembled complex, in contrast to earlier topology predictions (**Fig. 1A**). Hydrophobicity was calculated using Chimera. **(F)** Consensus predicted TM helices painted on to one copy each of P, Q and R (colored as in **Fig. 1A**).

Dissecting the complex further reveals that the subunits are arranged in a right-handed helix (**Fig. 3A**). The structural equivalence of a P/Q pair to R (**Fig. 2C**) means that the structure is effectively 6 copies of an R-like object forming a single helical turn, with the 5 copies of P further adorned by the inserted “periplasmic domain” (**Fig. 3B**). Analysis of the helical parameters that relate subunits reveals differences between the base and the tip of the structure (**Fig. 3C**); the helical pitch is tighter at the top than at the bottom of the complex but with average values consistent with those previously determined for both flagellar rod/hook/filament ^12-14^ and injectisome needle assemblies ^15^. Like the more peripheral axial assemblies, the PQR complex is constructed from pairs of helices arranged into a spiral (**Fig. S5**), although the orientation and arrangement of the helical hairpins is distinct. Analysis of the electrostatic surface of the PQR complex reveals that the inner surface, which would be predicted to form the export channel, is positively charged (**Fig. S5**), a feature that is also shared by the known rod/hook/flagellum/needle structures. A further important observation is that the PQR complex is seen to be closed at multiple points within the helical assembly. The base is closed by the highly conserved central portion of Q (**Fig. 3D**, **Fig. S6**). Immediately beyond this is a further closure formed by the highly conserved Met-Met-Met loop of P and finally a 15 residue insertion at the equivalent position in R forms a plug. Mutation of this insertion can overcome secretion defects caused by mutations in other components of the EA ^16^. Above these closure points is a small lumen that is then sealed at the top end by the N-terminal helices of P and R (**Fig. 3D**).

**Fig. 3.**
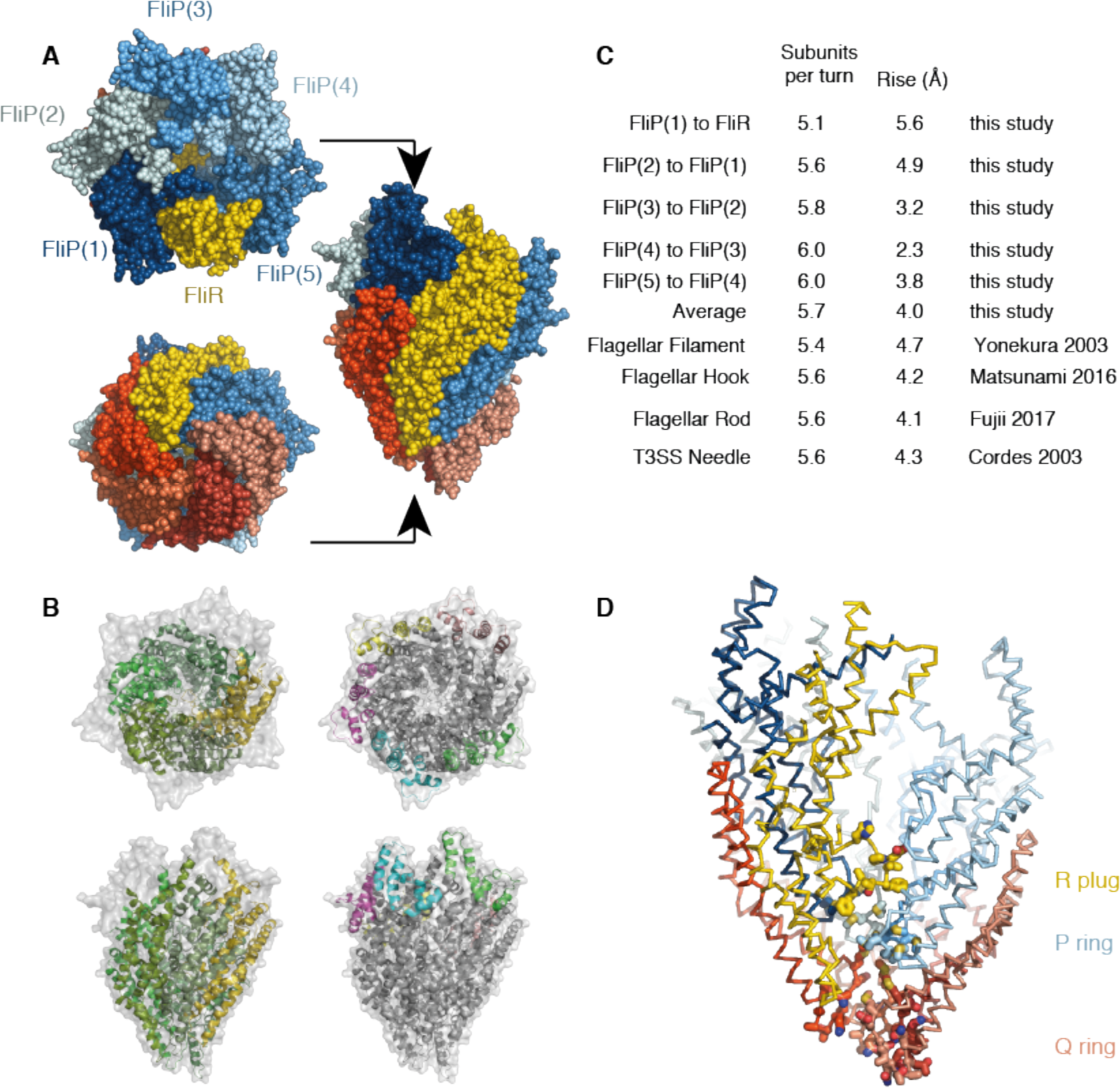
The P_5_Q_4_R_1_ complex is a right-handed helical assembly with helical parameters consistent with flagellar and injectisome assemblies. **(A)** Space filling representation of the P_5_Q_4_R_1_ complex with R (yellow), P (shades of blue) and Q (shades of red). **(B)** Surface representation of the intact P_5_Q_4_R_1_ complex (grey) containing a cartoon representation colored to highlight the helical rise. The left hand panel shows the core fold common to the P/Q heterodimers and R (left), while the right hand panel shows the “periplasmic domain” inserted into the P sequence. **(C)** Helical parameters calculated from transitioning between subunit pairs (labelled as in panel A). For comparison, previously determined for flagellar filament/hook and injectisome needle parameters are shown. **(D)** A backbone trace of the complex (colored as in **(A)**) with one copy of P removed to allow visualization of the interior. This reveals the central void is closed at the bottom by the layered “plug” loop in R and the highly conserved P and Q loops (highlighted with stick side chains).

The overall dimensions, helicity and hydrophobicity patterning of the complex do not support a standard localization within the inner membrane. We therefore sought to determine the location of the PQR complex in the assembled T3SS. Inspection of earlier single particle reconstructions of isolated basal bodies ^17^ and *in vivo* tomograms from both flagellar ^18^ and injectisome ^19^ T3SSs demonstrated that the PQR complex forms the structure previously described as the rod “cup and socket” ^8,20^. This is particularly striking when fitting our model within the highest resolution structure of a basal body determined to date, where the height, diameter and shape of the density seen in the basal body are an excellent match (**Fig. 4A**) ^17^. This therefore suggests that, in this reconstruction of a rod/needle-less system, the PQR complex is in the closed conformation that we observe. Placing the helical PQR complex in the cup/socket location predicts close contacts between it and the circularly symmetric components (FliF and SctC/SctJ in flagellar and injectisome systems respectively). Using *in vivo* photo-crosslinking and chemical crosslinking-mass spectrometry in the *Salmonella* injectisome system we were able to detect two residues in P on the outer surface of our complex that cross-linked to SctC (SpaP K132 and K135 to InvG K38) and a residue in SctJ (E138pBpa) that crosslinks to both P and R, providing further support for this location of the complex (**Fig. 4B**, **Fig. S7-10**).

**Fig. 4.**
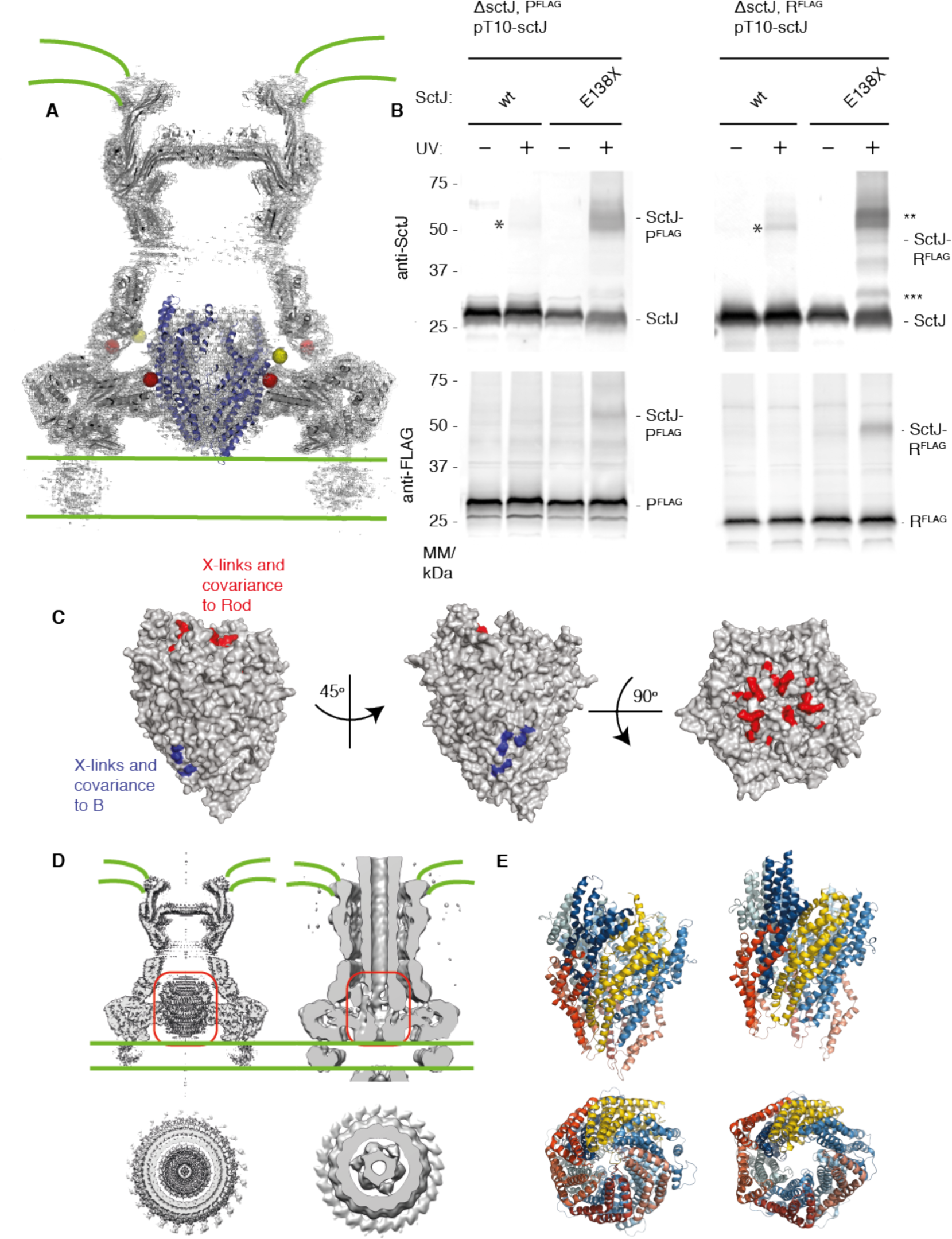
The P_5_Q_4_R_1_ complex is a core component of the basal body and forms a platform for assembly of the Rod. **(A)** Positioning our structure within an earlier high resolution reconstruction of the injectisome basal body ^17^ (grey surface and cartoon) reveals that the P_5_Q_4_R_1_ complex (blue cartoon) fits the un-occupied density in the center of the basal body. This region of the basal body has previously been called the “cup and socket” and sits above the proposed inner membrane location (shown as green lines). Residues on P that can be crosslinked to the basal body are highlighted by yellow spheres at the Ca position, while residues on SctC and SctJ of the basal body that cross-link to the PQR complex are shown in red. **(B)** *In vivo* photo-crosslinking studies reveal cross-links between P_5_Q_4_R_1_ and the inner membrane ring component SctJ in the *Salmonella* injectisome. The residues involved are highlighted in **(A)**. _*_SctJ-SctJ cross-link, tends to cross-link at a low level in the absence of pBpa. _**_indicates a band likely arising from the abundant SctJ-P crosslink. _***_indicates a band likely resulting from a SctJ-R crosslink. **(C)** Mapping of earlier cross-linking and our co-variation data (**Table S3**) onto P_5_Q_4_R_1_ reveal probable binding sites for B (FlhB/SctU) and the (inner) rod components. **(D)** Earlier basal body reconstructions show that the P_5_Q_4_R_1_ complex is closed in the absence of inner rod components (LHS, ^17^) and open in their presence (RHS, boxed in red, ^20^) **(E)** the open state of the P_5_Q_4_R_1_ complex can be modelled (RHS) by superposition of Rosetta co-variation based models for P and R onto our structure of the closed (LHS) complex with no clashes needing to be resolved.

The socket location of the complex suggests that the rod, which has previously been suggested to assemble directly onto the annular FliF ring ^18,21^ in the flagellar system, will actually assemble onto the already helical PQR complex. In order to investigate this further, we mapped the location of residues on PQR that have been shown to cross-link to the rod proteins ^9^. We also identified residues that strongly co-evolve between PQR and rod subunits (**Table S3**) ^11^. All of these amino acids map to the N-terminal helices on P and R and to the extreme C-termini of the rod proteins (including the Val 99 residue mutant of FliE that drastically reduces flagellation when mutated ^21^). This places the rod binding site at the top of the socket/EA complex, inside the walls formed from the “periplasmic domains” of the P subunits (**Fig. 4C**). Our structure therefore answers a long posited question of how the flagellum/needle helix nucleates; the asymmetric 5+1 structure drives the formation of the helical pitch.

Another consequence of the socket localization of the PQR portion of the export apparatus is that it places it above the predicted inner membrane position (**Fig. 4A**). Only the very tip of the Q-ring would be in contact with a predicted bilayer, despite the clear hydrophobic band that extends around the outer surface of the Q-subunits (**Fig. 2E**). Two possibilities to explain this observation are that either the bilayer inside the basal body is massively distorted, and/or these hydrophobic patches are covered by further protein interactions with the other two putative integral membrane proteins, A (FlhA/SctV) & B (FlhB/SctU). The A protein forms a circular, nonameric complex with a large domain of known structure localized in the cytoplasm immediately below the basal body ^22,23^. The integral membrane domain of this protein would therefore be predicted to lie in the inner membrane around the base of our PQR complex. We note that *in situ* tomograms of the *Salmonella* injectisome show a distinct density in this location but that cells deficient in A not only lack this density but also demonstrate a distorted membrane leaflet that reaches up to the base of the Q subunits ^19^ (**Fig. S11**). However, evidence for direct association via cross-linking or co-evolution is lacking, and production of stable PQRA complexes has proved challenging ^22^. By contrast, both co-variance (**Table S3**) ^11^ and *in vivo* photo cross-linking studies ^9^ imply specific and direct interactions between PQR and B. Mapping the residues implicated in the B interaction onto our structure suggests it assembles at the interface between R and the Q-less P at the base of our helical assembly (**Fig. 4C**).

The fact that the isolated PQR complex adopts a closed conformation is likely important for maintaining bacterial viability, by ensuring the complex does not form breaches in the cell membrane prior to full assembly of the basal body and control of gating by the entire T3SS. As noted above, the shape of our closed complex is consistent with the shape of the socket/cup seen in a rod-less basal body and this, coupled with the relatively tight fit of the complex against the FliF/SctJ ring, suggests that the initially assembled basal body is impermeable to substrate. However, we noted that the shape of the socket/cup density in reconstructions of more intact basal bodies containing needles ^20,24^ was more open at the base of the density now ascribed to PQR (**Fig. 4D**). We therefore sought to model this open state by allowing the conformation of our P and R subunits to relax back to the much straighter arrangement of helices found in the models derived purely from co-evolution data. This had the effect of opening the closures at the base of the complex via a concerted iris mechanism, producing a model without significant atomic clashes (**Fig. 4E** and **Movie S2**). Such a conformation would therefore allow substrate access to a secretion channel through the centre of the PQR complex and, given the cross-linking and co-evolution data, permit assembly of the rod on the top of the export gate.

This study has demonstrated the following regarding the PQR export gate complex: 1) that it has a core stoichiometry that is conserved across flagellar and injectisome T3SS; 2) that it forms a structure with an unpredicted helical symmetry; 3) that it sits above the likely location of the bacterial inner membrane as a core component of the basal body. The data suggest that the core export gate complex is contiguous with the helical axial components that culminate in the flagellum/needle, and therefore that the export pathway for secreted substrates will be through the center of this complex, directly into the channel within the rod and filament or needle. The next major question that therefore arises from this observation is that of the nature of the mechanism of initially opening the PQR channel for secretion and the role of the other EA components in this process. Subsequently, whether assembly of the rod onto the opened PQR complex is then sufficient to lock it in the open conformation, or whether it plays a role in further gating, will require investigation. Regardless, the central position and core role of this complex in the secretion pathway of many virulent bacteria make it an attractive target for future drug development, especially in light of increased antibiotic resistance.

## Acknowledgments

We thank Errin Johnson & Adam Costin of the Central Oxford Structural Microscopy and Imaging Centre and all staff of the ARC Centre of Advanced Imaging, Monash University, for assistance with data collection. Hans Elmlund (Monash) is thanked for assistance with data collection, processing strategies and access to SIMPLE code ahead of release. For collection of preliminary data we acknowledge Diamond for access and support of the Cryo-EM facilities at the UK national electron bio-imaging centre (eBIC), proposal EM15354, funded by the Wellcome Trust, MRC and BBSRC. We thank Thomas Marlovits for providing PrgK (SctJ) antisera. The Central Oxford Structural Microscopy and Imaging Centre is supported by the Wellcome Trust (201536), The EPA Cephalosporin Turst, the Wolfson Foundation and a Royal Society/Wolfson Foundation Laboratory Refurbishment Grant (WL160052). Work performed in the lab of S.M.L was supported by a Wellcome Trust Investigator Award (100298) and an MRC programme grant (M011984). Work performed in the lab of C.V.R. is funded by a Wellcome Trust Investigator Award (104633), an ERC Advanced Grant ENABLE (641317) and an MRC programme grant (N020413). J.G. is a Junior Research Fellow of The Queen’s College, Oxford. Work performed in the laboratory of S.W. was supported by the Alexander von Humboldt Foundation in the framework of the Sofja Kovalevskaja Award endowed by the Federal Ministry of Education and Research (BMBF) (www.avh.de) and by the Deutsche Forschungsgemeinschaft (DFG) as part of the Collaborative Research Center (SFB) 766 Bacterial cell envelope, project B14 (www.dfg.de).

## Author Contributions

All authors contributed to editing of manuscript and figures.

Lucas Kuhlen: Performed experiments. Strain and plasmid construction, complex purification, native mass spectrometry, cryoEM grid optimisation, cryoEM data analysis and model building and analysis.

Patrizia Abrusci: Performed experiments. Strain and plasmid construction, complex purification initial cryoEM grid optimisation and cryoEM data analysis.

Steven Johnson: Performed experiments. Designed and supervised experiments.

Characteristion of protein complexes. CryoEM data analysis, structure determination and analysis. Wrote first draft of manuscript and figures with SML.

Joseph Gault: Performed and supervised native mass spectrometry experiments.

Justin Deme: Performed experiments. CryoEM grid optimisation and data collection Joseph Caesar: Performed experiments. Optimised computational analysis of cryo-EM data. Tobias

Dietsche: Performed experiments. Strain and plasmid construction, *in vivo* photocrosslinking

Mehari Tesfazgi Mebrhatu: Performed experiments. NC purification, chemical crosslinking Tariq Ganief: Performed experiments. MS of DSS crosslinked samples. Analysed data.

Boris Macek: Analysed DSS MS data. Wrote paper: DSS MS methods section.

Samuel Wagner: Analysed data: *in vivo* photocrosslinking, chemical crosslinking, DSS MS. Wrote first draft of photo‐ and chemical-crosslinking sections.

Carol V Robinson: Supervised design and implementation of native mass spectrometry experiments

Susan M Lea: Performed and supervised experiments. CryoEM grid optimisation and data collection, data and structure analysis. Wrote first draft of paper with SJ.

## References and Notes

1 Abrusci, P., McDowell, M. A., Lea, S. M. & Johnson, S. Building a secreting nanomachine: a structural overview of the T3SS. Curr Opin Struct Biol 25, 111-117, doi:10.1016/j.sbi.2013.11.001 (2014).

2 Erhardt, M., Namba, K. & Hughes, K. T. Bacterial nanomachines: the flagellum and type III injectisome. Cold Spring Harb Perspect Biol 2, a000299, doi:10.1101/cshperspect.a000299 (2010).

3 Buttner, D. Protein export according to schedule: architecture, assembly, and regulation of type III secretion systems from plant‐ and animal-pathogenic bacteria. Microbiol Mol Biol Rev 76, 262-310, doi:10.1128/MMBR.05017-11 (2012).

4 Deng, W. et al. Assembly, structure, function and regulation of type III secretion systems. Nat Rev Microbiol 15, 323-337, doi:10.1038/nrmicro.2017.20 (2017).

5 Macnab, R. M. How bacteria assemble flagella. Annu Rev Microbiol 57, 77-100, doi:10.1146/annurev.micro.57.030502.090832 (2003).

6 Fabiani, F. D. et al. A flagellum-specific chaperone facilitates assembly of the core type III export apparatus of the bacterial flagellum. PLoS Biol 15, e2002267, doi:10.1371/journal.pbio.2002267 (2017).

7 Fukumura, T. et al. Assembly and stoichiometry of the core structure of the bacterial flagellar type III export gate complex. PLoS Biol 15, e2002281, doi:10.1371/journal.pbio.2002281 (2017).

8 Wagner, S. et al. Organization and coordinated assembly of the type III secretion export apparatus. Proc Natl Acad Sci U S A 107, 17745-17750, doi:10.1073/pnas.1008053107 (2010).

9 Dietsche, T. et al. Structural and Functional Characterization of the Bacterial Type III Secretion Export Apparatus. PLoS Pathog 12, e1006071, doi:10.1371/journal.ppat.1006071 (2016).

10 Zilkenat, S. et al. Determination of the Stoichiometry of the Complete Bacterial Type III Secretion Needle Complex Using a Combined Quantitative Proteomic Approach. Mol Cell Proteomics 15, 1598-1609, doi:10.1074/mcp.M115.056598 (2016).

11 Ovchinnikov, S., Kamisetty, H. & Baker, D. Robust and accurate prediction of residue-residue interactions across protein interfaces using evolutionary information. Elife 3, e02030, doi:10.7554/eLife.02030 (2014).

12 Yonekura, K., Maki-Yonekura, S. & Namba, K. Complete atomic model of the bacterial flagellar filament by electron cryomicroscopy. Nature 424, 643-650, doi:10.1038/nature01830 (2003).

13 Fujii, T. et al. Identical folds used for distinct mechanical functions of the bacterial flagellar rod and hook. Nat Commun 8, 14276, doi:10.1038/ncomms14276 (2017).

14 Matsunami, H., Barker, C. S., Yoon, Y. H., Wolf, M. & Samatey, F. A. Complete structure of the bacterial flagellar hook reveals extensive set of stabilizing interactions. Nat Commun 7, 13425, doi:10.1038/ncomms13425 (2016).

15 Cordes, F. S. et al. Helical structure of the needle of the type III secretion system of Shigella flexneri. J Biol Chem 278, 17103-17107, doi:10.1074/jbc.M300091200 (2003).

16 Hara, N., Namba, K. & Minamino, T. Genetic characterization of conserved charged residues in the bacterial flagellar type III export protein FlhA. PLoS One 6, e22417, doi:10.1371/journal.pone.0022417 (2011).

17 Worrall, L. J. et al. Near-atomic-resolution cryo-EM analysis of the Salmonella T3S injectisome basal body. Nature, doi:10.1038/nature20576 (2016).

18 Zhao, X. et al. Cryoelectron tomography reveals the sequential assembly of bacterial flagella in Borrelia burgdorferi. Proc Natl Acad Sci U S A 110, 14390-14395, doi:10.1073/pnas.1308306110 (2013).

19 Hu, B., Lara-Tejero, M.., Kong, Q., Galan, J. E. & Liu, J. In Situ Molecular Architecture of the Salmonella Type III Secretion Machine. Cell 168, 1065-1074e1010, doi:10.1016/j.cell.2017.02.022 (2017).

20 Schraidt, O. & Marlovits, T. C. Three-dimensional model of Salmonella’s needle complex at subnanometer resolution. Science 331, 1192-1195, doi:10.1126/science.1199358 (2011).

21 Minamino, T., Yamaguchi, S. & Macnab, R. M. Interaction between FliE and FlgB, a proximal rod component of the flagellar basal body of Salmonella. J Bacteriol 182, 30293036 (2000).

22 Abrusci, P. et al. Architecture of the major component of the type III secretion system export apparatus. Nat Struct Mol Biol 20, 99-104, doi:10.1038/nsmb.2452 (2013).

23 Chen, S. et al. Structural diversity of bacterial flagellar motors. EMBO J 30, 2972-2981, doi:10.1038/emboj.2011.186 (2011).

24 Radics, J., Konigsmaier, L. & Marlovits, T. C. Structure of a pathogenic type 3 secretion system in action. Nat Struct Mol Biol 21, 82-87, doi:10.1038/nsmb.2722 (2014).

25 Ashkenazy, H. et al. ConSurf 2016: an improved methodology to estimate and visualize evolutionary conservation in macromolecules. Nucleic Acids Res 44, W344-350, doi:10.1093/nar/gkw408 (2016).

